# A computational account of joint SSRI and anti-inflammatory treatment

**DOI:** 10.1101/2023.12.26.573389

**Authors:** Melissa Reneaux, Helen Mayberg, Karl Friston, Dimitris A. Pinotsis

**Affiliations:** Centre for Mathematical Neuroscience and Psychology and Department of Psychology, City —University of London, London EC1V 0HB, United Kingdom; Psychology and Behaviour Program, School of Liberal Studies, UPES, Dehradun 248007, India; Icahn School of Medicine at Mt. Sinai, New York, 10029, NY, USA; Wellcome Centre for Human Neuroimaging, UCL, London WC1N 3AR, United Kingdom; The Picower Institute for Learning & Memory and Department of Brain and Cognitive Sciences, Massachusetts Institute of Technology, Cambridge, MA 02139, USA

**Keywords:** Immune system, inflammation, pro-inflammatory cytokines, serotonin, glutamate, prefrontal cortex, subcallosal cingulate cortex

## Abstract

We present a computational model that elucidates the interplay between inflammation, serotonin levels, and brain activity. The model delineates how inflammation impacts extracellular serotonin, while cerebral activity reciprocally influences serotonin concentration. Understanding the reciprocal interplay between the immune system and brain dynamics is important, as unabated inflammation can lead to relapsing depression. The model predicts dynamics within the prefrontal cortex (PFC) and subcallosal cingulate cortex (SCC), mirroring patterns observed in depressive conditions. It also accommodates pharmaceutical interventions that encompass anti-inflammatory and antidepressant agents, concurrently evaluating their efficacy with regard to the severity of depressive symptoms.

## Introduction

Depression affects 4.4% of the world’s population [WHO 2017; Malhi & Mann 2018]. Putative causes are multifactorial. They include monoamine depletion, anxiety, stress and inflammation, all of which mutually interact. These factors affect activity in distinct brain circuits regulating emotional behaviour, including the dorsal raphe nucleus (DRN) and other infralimbic and prelimbic cortices and frontal areas [Sarter *et al*. 1984]. Depression is thus considered a circuit disorder [Mayberg 2007; Dunlop *et al*. 2023; Gray *et al*. 2023]. Studies have revealed the central role of circuits like the subcallosal cingulate cortex (SCC) [Mayberg *et al*. 1997; Mayberg *et al*. 1999; Greicius *et al*. 2007], default mode network [Dunlop *et al*. 2023] and executive control network [Liu *et al*. 2020].

Depression is a chronic disorder. A large number of patients with depression relapse [Touya *et al*. 2022]. Recurrent depression and disease progression are common [Kessing & Andersen 2017]. Treatments work often but not always [Gabriel *et al*. 2020]. Some patients develop treatment non-response or resistance [Voineskos *et al*. 2020]. One cause underlying disease chronicity and non-response is inflammation [Miller *et al*. 2009; Miller & Timmie 2009; Dantzer *et al*. 2008]. Inflammation has profound effects on interoception and ensuing affective regulation [McEwen & Gianaros 2011; Peters *et al*. 2017]. It constitutes a threat to allostasis, i.e., the ability of the brain to resolve and pre-empt environmental and internal challenges to the body by adapting physiological parameters [McEwen 1998; Cohen *et al*. 2007]. About a quarter of depressed patients exhibit inflammation [Osimo *et al*. 2019]. Some patients show resistance to anti-depressants but respond to anti-inflammatory drugs [Carvalho *et al*. 2013; Yoshimura *et al*. 2009; Hodes *et al*. 2015]. In brief, depression is a circuit illness affected severely by inflammation. To understand its pathology and the link between inflammation and depression we developed a computational model. Previous work with similar models have described alterations in excitation to inhibition balance [Page & Coutellier 2019], the role of serotonin and other monoamines in depression [Kringelbach *et al*. 2020; Ramirez-Mahaluf *et al*. 2017] and changes in the HPA axis [Pariante & Lightman 2008]. From a purely theoretical standpoint, [Bhat *et al*. 2021] consider the complementary alterations of immunological sensitivity as an analogue of sensory attenuation.

Here, we focused on interactions between the immune system, the serotonergic system and brain activity. We modelled neuronal responses to TNFα concentration changes. TNFα is the acronym of a common cytokine known as Tumor Necrosis Factor alpha [Raison *et al*. 2013; Yao *et al*. 2020; Brymer *et al*. 2019]. Cytokines are a key part of the immune system: they are chemicals used by immune cells for communication. They are elevated in inflammation and have altered levels in inflammation and depression [Dowlati *et al*. 2010; Köhler *et al*. 2017].

Specifically, we modelled how TNFα affects neural activity in the prefrontal cortex (PFC) and the subcallosal cingulate gyrus (SCC). We focused on these areas because they show highly consistent depression-related abnormalities and are the main targets for treatment [Fox *et al*. 2012]. Our model includes excitatory and inhibitory neuronal populations driven by NMDA, GABA and serotonergic currents, which evince the dynamics of 5HT1A and 5HT2A receptors. The amplitude of serotonergic currents was determined by extracellular serotonin concentration. In turn, this was modulated by changes in serotonin synthesis and reuptake due to inflammation. We quantified the impact of inflammation using the ratio of TNFα concentration in patients vs. control — that we called degree of inflammation [Zou *et al*. 2018]. Finally, we introduced expressions that link this degree to serotonin synthesis and reuptake.

Technically, our model is a neural mass model based upon a set of stochastic differential equations describing the kinetics of serotonin and synaptic dynamics of coupled neuronal populations. Neuronal activity corresponds to the mean synaptic activity (modelled by synaptic gating variables, equipped with random fluctuations) at steady-state. Numerically, neuronal activity is quantified by the mean (and standard error of the mean: sem) over multiple solutions of the differential equations, for any given set of their parameters. The ensuing neural mass model predicts the impact of peripheral inflammation on depression for different cytokine and serotonin concentrations. It also explains how a joint SSRI and anti-inflammatory treatment might ameliorate depression.

## Results

### A computational model of inflammation-mediated changes of serotonergic availability

Our model links inflammation, the serotonergic system and brain activity. It describes how inflammation alters serotonergic currents and ensuing brain activity. The effect of inflammation on serotonin is two-fold. This is shown in Figures 1A and 1B, see also [Liu *et al*. 2020; Dowlati *et al*. 2010]. (i) Reduction of serotonin synthesis: cytokines stimulate the production of an enzyme called indoleamine 2,3-dioxygenase (IDO). This directs tryptophan, a precursor of serotonin, into the kynurenine pathway: tryptophan produces more kynurenine and other metabolites, instead of serotonin [Miller *et al*. 2013]. (ii) Increase of serotonin reuptake: the enzyme IDO increases the activity of serotonin transporters. These are proteins that clear serotonin from the extracellular medium by transporting it back into the presynaptic terminal [Miller *et al*. 2009].

**Figure 1:**
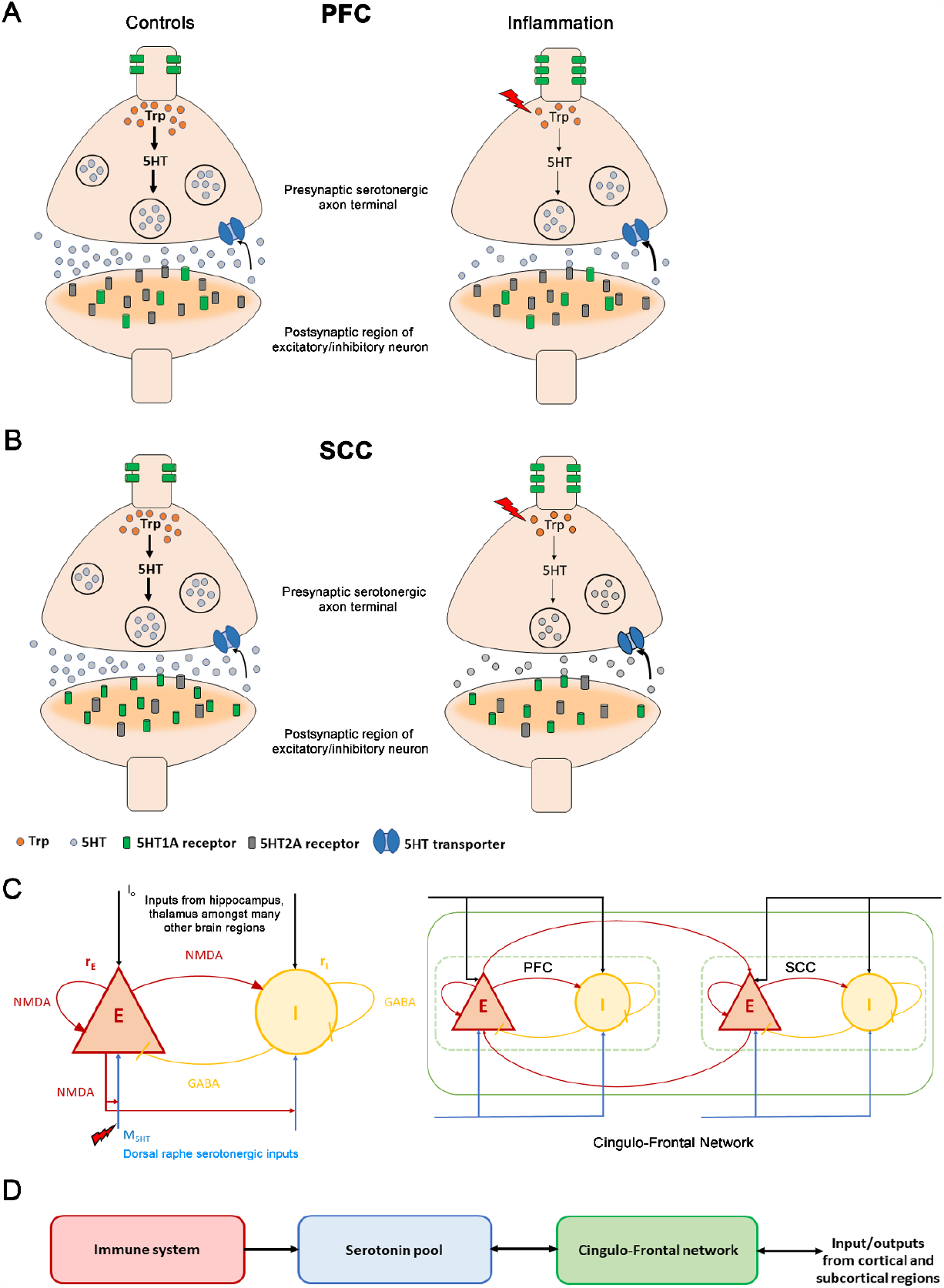
The immune-brain interaction. **[A]** Changes between control and inflammation at a synaptic terminal in the prefrontal cortex (PFC). In control subjects (left panel), tryptophan (orange discs), a precursor of serotonin, is converted to serotonin (grey discs) in the presynaptic serotonergic neuron. Serotonin is then packaged in synaptic vesicles (large black circles). Synaptic vesicles move towards the synaptic bouton and fuse to the membrane releasing serotonin into the extracellular medium. The extracellular serotonin binds to the postsynaptic receptors (grey and green cylindrical structures) and signals a cascade of events in the postsynaptic excitatory/ inhibitory neuronal terminal. The excitatory serotonergic 5HT2A receptors (grey) are abundantly present in PFC compared to inhibitory 5HT1A receptors (green). 5HT1A auto-receptors are located on the somato-dendritic regions of serotonergic neurons and regulate serotonergic neuronal activity. The unused extracellular serotonin is re-uptaken (black arrow) by presynaptic serotonin transporters (blue channel), to be recycled and reused. During inflammation (right panel, red lightning bolt), TNFα concentration increases (red lightning bolt). This reduces tryptophan (fewer orange discs in right vs. left hand panel) involved in serotonin synthesis (thinner black vertical arrows). Also, it leads to fewer synaptic vesicles containing serotonin (fewer black circles in presynaptic neuron). Less serotonin is released. Inflammation also causes serotonin transporters to rapidly reuptake the available extracellular serotonin (bold black arrow). Further, an increase in 5HT1A auto-receptors is observed. **[B]** Inflammation changes at a synaptic terminal in the subcallosal cingulate cortex (SCC). The postsynaptic receptor distribution is complementary to that of the PFC: the SCC has an abundance of inhibitory 5HT1A receptors (green cylinders) compared to the excitatory 5HT2A receptors (grey cylinders). Under inflammation (right panel), a reduction in serotonin is observed. In depression, a reduction in postsynaptic 5HT1A receptors and an increase in the 5HT1A auto-receptors is observed. **[C]** (Left panel) Excitatory (red) and inhibitory (yellow) neuronal populations and their connections. Excitatory currents are mediated by NMDA receptors. Inhibitory by GABA receptors. Brain activity is coupled with the neurotransmitter system. Io (black arrows) are external currents received from other cortical and subcortical brain regions. Serotonergic currents, *M*5*HT* are shown by blue arrows. Inflammation causes a *M*5*HT* reduction. Right panel: PFC-SCC brain network. The two regions are coupled by long-range NMDA connections targeting excitatory populations. **[D]** Elevated TNFα concentration (red module) due to inflammation reduces serotonin (blue module) by impacting synthesis and reuptake. This, in turn, reduces serotonergic input to the PFC-SCC network (green module). Excitatory population activity within PFC and SCC reciprocally modulates the serotonin concentration (double arrow). Further, the PFC-SCC circuit receives — and sends — signals to many other regions in the brain, like hippocampus, thalamus and amygdala.

Several cytokines including Interleukin 6 (IL6), C-reactive protein (CRP) and TNFα have elevated levels in depression [Miller *et al*. 2009]. TNFα is the most relevant for depression as it regulates extracellular serotonin levels [Ma *et al*. 2016; Hestad *et al*. 2003; Zou *et al*. 2018]. Peripheral administration of TNFα antagonist has been shown to improve depressive mood [Tyring *et al*. 2006], reduce fatigue [Monk *et al*. 2006] and alleviate depression [Persoons *et al*. 2005]. Furthermore, TNFα receptor knock out mice show reduced anxiety-like behaviour during immune activation [Silverman *et al*. 2007].

Our model describes how TNFα concentration affects brain activity; specifically, in the cingulo-frontal circuit thought to underlie depression (Figure 1). It rests on the premise that depression is a circuit disorder [Mulders *et al*. 2015; Mayberg 2007; Greicius *et al*. 2007] affecting the frontal cortex, insula, thalamus and other areas. Among them, the circuit modelled here shows highly consistent depression-related abnormalities and is the target of treatments including medication, psilocybin treatments and psychotherapy [Fox *et al*. 2012]. This circuit includes PFC and SCC is also the target of Deep brain stimulation (DBS) [Mayberg 1999]. Transcranial magnetic stimulation (TMS) impacts this circuit— changing the balance of neural activity between and finding the spot with maximal anti-correlation of the SCC and PFC [Liston *et al*. 2014].

To our knowledge, the model described in this foundational paper is the first to address how changes in cytokine concentration affect neural activity. The neural mass model includes excitatory and inhibitory neuronal populations driven by excitatory NMDA currents (shown in red, Figure 1C, left panel), inhibitory GABA currents (yellow) and external input *I*_*o*_ (black), see also [Ramirez-Mahaluf *et al*. 2017] who described glutamate dysregulation and its effects on treatment response and EEG rhythms. Additionally, our model includes serotonergic currents *M*_5*HT*_ (blue), whose amplitude is determined by the extracellular serotonin concentration. This concentration is modulated by changes in serotonin synthesis and reuptake due to inflammation (*Methods*) [Kringelbach *et al*. 2020]. A novel feature of our model is the dependence of serotonin synthesis and reuptake on changes in cytokine concentration due to elevated TNFα levels of the sort observed in inflammation. Below we explain this dependence in detail.

Serotonin synthesis (release) depends on the density of the fibres connecting the DRN with SCC or PFC. It also depends on the activity of the excitatory neuronal population in SCC or PFC. The excitatory long-range reciprocal interaction between the PFC and SCC is depicted in Figure 1C (right panel). In general, receptor expression varies systematically across cortical areas [Froudist-Walsh *et al*. 2023]. Among all receptors, serotonin shares the most marked change (gradient) in expression over the cortex. This is also the case in the regions considered here. PFC has an abundance of excitatory 5HT2A receptors (Figure 1A), while SCC has inhibitory 5HT1A receptors [Figure 1B; Palomero-Gallagher *et al*. 2009]. Both the brain regions have 5HT1A and 5HT2A serotonergic receptors. The receptor densities of 5HT1A in the PFC and 5HT2A in the SCC are low (Figure 1A, B). Thus, we modelled contributions of 5HT2A receptors in PFC and 5HT1A receptors in SCC. Model parameters — expressing receptor densities — were based on measurements with positron emission tomography (PET) [Kringelbach *et al*. 2020]. Inflammation parameters were chosen to obtain serotonin levels reported in depression studies. These and other model parameters can be found in Tables 1 and 2. They follow [Kringelbach *et al*. 2020; Zou *et al*. 2018]. Elevated levels of TNFα (red box in Figure 1D) cause a reduced synthesis and increased reuptake of extracellular serotonin (blue box). This, in turn, alters activity in the cingulo-frontal brain network (green box in Figure 1D).

**Table 1:**
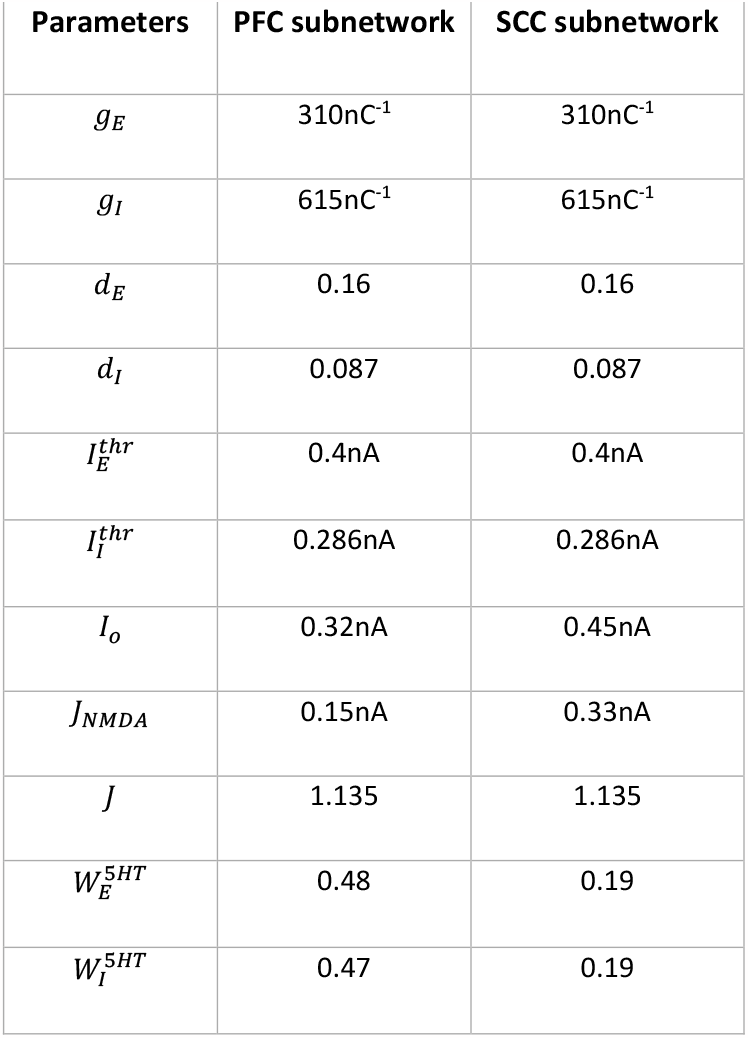
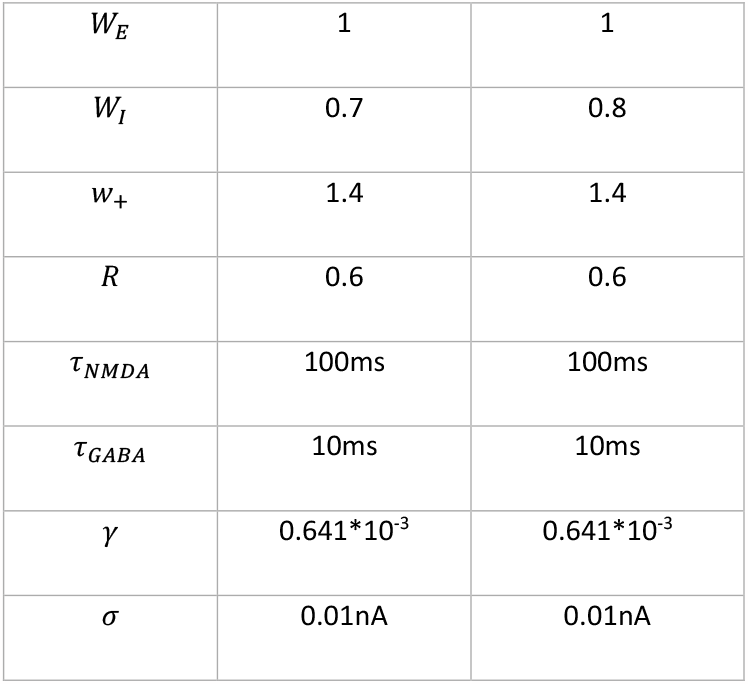
Neuronal model parameters.

**Table 2:** List of parameters for serotonergic inputs.

| Parameters | PFC subnetwork | SCC subnetwork |
| --- | --- | --- |
| $\alpha$ | 5 | 5 |
| $C_{BR}$ | 15 | 15 |
| $V_{max}$ | $1300\text{nMs}^{-1}$ | $1300\text{nMs}^{-1}$ |
| $K_m$ | 170 (Control condition)<br>200 (SSRI treatment) | 170 (Control condition)<br>200 (SSRI treatment) |
| $c_1$ | 5 | 4 |
| $c_2$ | 0.019 | 0.019 |
| $m$ | 2 | 2 |
| $\tau_{5HT}$ | 120ms | 120ms |
| $J_{5HT}$ | 1 | 1 |
| $\beta$ | 0.008 | 0.008 |
| $[Cyt]/[Cyt]_b$ | 1 (Control condition)<br>1.25 (Mild inflammation)<br>1.4 (Moderate inflammation)<br>2.3 (Severe inflammation) | 1 (Control condition)<br>1.25 (Mild inflammation)<br>1.4 (Moderate inflammation)<br>2.3 (Severe inflammation) |
| $B$ | 0.55 (Anti-inflammatory treatment) | 0.55 (Anti-inflammatory treatment) |

### Inflammation leads to cytokine increase and reduced serotonin

Inflammation increases the concentration of the cytokine TNFα. We first asked how this affects extracellular serotonin levels in PFC and SCC. TNFα increase stimulates the enzyme IDO that drives tryptophan into producing kynurenine and impairs serotonin synthesis. IDO also increases serotonin reuptake. Our model parametrizes the change in TNFα concentration as a result of inflammation (see *Methods* and below). It also predicts the corresponding changes in serotonin synthesis and reuptake that result from TNFα changes.

We characterised the increase in TNFα as the ratio [*Cyt*]/[*Cyt*]_*b*_. We call this, the degree of inflammation. It describes the relative increase in the TNFα cytokine concentration [*Cyt*] relative to controls, denoted by [*Cyt*]_*b*_. [*Cyt*]_*b*_ is equal to 2.69±0.14pg/mL (mean ± sem) as measured in human serum [Zou *et al*. 2018, Figure 2A]. We considered a graded rise in the degree of inflammation. Our analysis followed [Zou *et al*. 2018]. These authors used the Hamilton Depression Rating Scale (HAMD) scores and found linear correlations between depression severity and TNFα concentration. Similar to Zou *et al*. [Zou *et al*. 2018], we considered three inflammation conditions: mild, moderate and severe (Figure 2B). A 1.25-fold rise in inflammation degree corresponds to mild inflammation where [*Cyt*] was 3.35±0.13pg/mL. A 1.4-fold rise corresponds to moderate inflammation and a 2.3-fold rise to severe inflammation. The corresponding cytokine concentrations [*Cyt*] were 3.79±0.08pg/mL and 6.19±0.72pg/mL respectively [Zou *et al*. 2018, Figure 2A, two rightmost bars].

**Figure 2:**
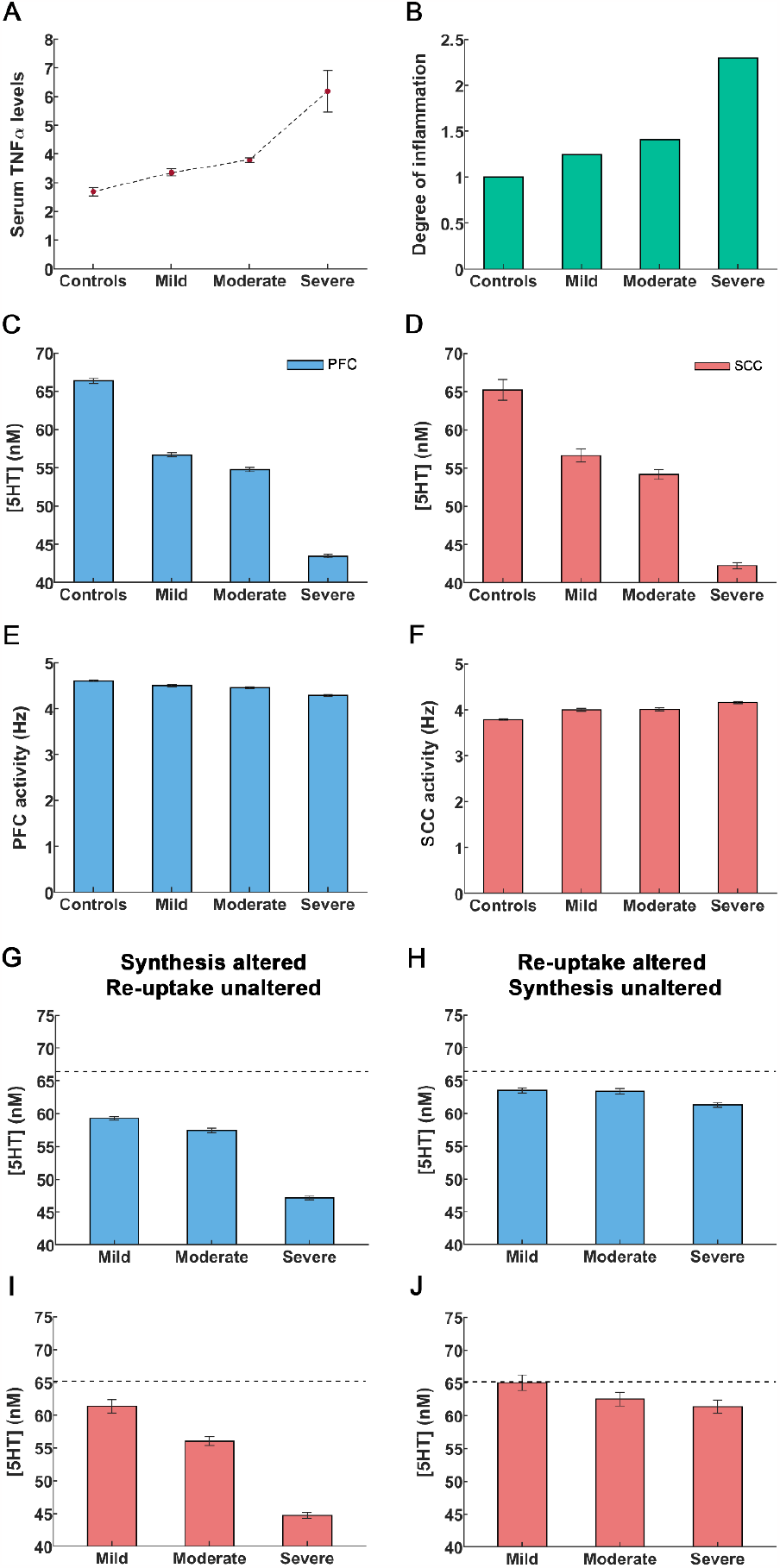
An increase in cytokines caused by peripheral inflammation leads to reduction in serotonin concentration and alterations in cingulo-frontal network activity. **[A]** Cytokine TNFα concentrations in controls are denoted by [*Cyt*]*b*. Concentrations in depressed individuals by [*Cyt*]. The solid circles refer to mean TNFα concentrations and errors bars are the sem values taken from [Zou *et al*. 2018]. **[B]** Using concentrations from [A], we constructed the concentration ratio [*Cyt*]/[*Cyt*]*b*.This is referred to as the degree of inflammation. The control condition corresponds to a degree of inflammation of one (leftmost bar). Other bars denote the degree corresponding to mild, moderate and severe inflammation as defined in [Zou *et al*. 2018]. **[C**,**D]** Rise in inflammation degree reduces serotonin concentrations in PFC and SCC. **[E**,**F]** Rise in inflammation degree reduces theta band activity. **[G**,**I]** The extracellular serotonin levels in the PFC and SCC assuming inflammation affects serotonin synthesis only (cf. *X1* term in Equation 10 in *Methods*) and reuptake is unaltered. **[H**,**J]** Extracellular serotonin levels assuming inflammation affects serotonin reuptake only (cf. *X2* term in Equation 10) and synthesis is unaltered. The black dash line in the figures G-I refer to the extracellular serotonin levels in controls. The error bars refer to the sem values obtained from 100 simulations run for each condition.

Our model, predicts the extracellular serotonin concentration for different degrees of inflammation. Serotonin concentration for controls 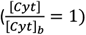 was predicted to be 66.35±0.35nM in PFC (Figure 2C, leftmost bar) and 65.21±1.32nM in SCC (Figure 2D, leftmost bar). These concentrations are similar to those reported in [Hersey *et al*. 2021]^1^. In severe depression, a 2.3-fold increase in the degree of inflammation led to a serotonin concentration of 43.48±0.22nM in PFC and 42.25±0.40nM in SCC [Hersey *et al*. 2021] (Figures 2C,D, rightmost bars). In mild inflammation concentration was 56.69±0.29nM in PFC and 56.63±0.82nM in SCC. Last, a 1.4-fold rise in the degree of inflammation (moderate) rendered the concentration 54.77±0.28nM in PFC and 54.16±0.66nM in SCC (Figures 2C,D). Serotonin concentrations for the control and severe inflammation conditions are similar to [Hersey *et al*. 2021]. Our model also predicted concentrations for mild and moderate depression considered in [Zou *et al*. 2018]. These are novel results. To the best of our knowledge no experimental data exists for these conditions.

Next, we distinguished inflammation effects on serotonin synthesis and reuptake. These were quantified by the *X*_*1*_ and *X*_*2*_ terms in our model (Equations 10 in *Methods*). Reduced synthesis was described by the *X*_*1*_ term, while elevated reuptake rates entailed by the *X*_*2*_ term. These terms capture differences due to excitatory 5HT2A receptors in PFC and inhibitory 5HT1A receptors in SCC. We then considered the effect of changing one of these terms and keeping the other constant. Assuming serotonin synthesis was altered without TNFα affecting reuptake (i.e., the *X1* term) led to a significant reduction in the extracellular serotonin levels (Figures 2G,I). These reductions were similar to those obtained above when assuming that TNFα affected both synthesis and reuptake (Figures 2C,D). Assuming that only serotonin reuptake was impacted by TNFα (*X2* term) led to a slight reduction in serotonin levels (Figures 2H,J). Thus, changes in serotonin concentration due to inflammation seem to be driven by TNFα effects on synthesis not reuptake.

### Dysregulation in cingulo-frontal network activity as a result of inflammation

We then turned to brain activity. Our model generates neuronal activity in PFC and SCC. Both are main hubs in a medial prefrontal network known to mediate chronic stress and depression [Dum *et al*. 2016]. We considered control responses and responses for different degrees of inflammation as above. For controls, resting state PFC activity was 4.61±0.02Hz (Figure 2E, leftmost bar), similar to observed monkey recordings [Wilson *et al*. 1994; Kim & Sejnowski 2021]. The model predicted reduced PFC neural activity for mild and moderate inflammation i.e. 4.50±0.02Hz and 4.46±0.02Hz respectively (Figure 2E, middle bars). Under severe inflammation, activity was reduced to 4.29±0.02Hz. This amounts to about 7% reduction in activity between control and severe inflammation conditions (Figure 2E, rightmost bar), similar to recordings by [Fernández-Palleiro *et al*. 2020; Fitzgerald *et al*. 2008].

Our model also predicted that SCC theta band activity increased when the degree of inflammation increased. Activity in controls was found to peak at 3.79±0.02Hz (Figure 2F, leftmost bar). Similar predictions for mild and moderate inflammation included peaks at 4.00±0.03Hz and 4.01±0.03Hz respectively (Figure 2F, middle bars). For severe inflammation, activity was found to peak at 4.16±0.02Hz (Figure 2F, rightmost bar). This amounts to a 10% increase in SCC network activity compared to controls, similar to recordings by [Pizzagalli 2011, Mayberg *et al*. 2005; Wolff *et al*. 2019].

Overall, inflammation had opposite effects in PFC and SCC see also [Shao *et al*. 2018]. This is explained by the difference of serotonergic receptors in these regions. PFC has an abundance of excitatory 5HT2A receptors, while SCC has an abundance of inhibitory 5HT1A receptors. Thus, our model found that frontal activity was reduced with inflammation, while limbic activity increased. Crucially, it predicted subtle changes and their dependence on the degree of inflammation that matched experimental recordings.

### SSRIs alleviate serotonin deficiency under mild inflammation

Next, we considered drug treatments, and their limitations. We first modelled SSRI effects. We asked whether SSRIs could alleviate depression in the presence of inflammation. SSRIs are commonly used as the first line of treatment for depression. In our model, their administration (reuptake inhibition, red “X” in Figure 3A) is characterized using the Michaelis-Menten constant *K*_*m*_. Serotonin reuptake followed the Michaelis-Menten kinetics. The constant *K*_*m*_ describes the binding affinity (or likeness) of serotonin to its transporter. A large *K*_*m*_ corresponds to small affinity (denominator in Equation 10, see *Methods*). In the presence of a reuptake inhibitor the affinity of serotonin to its transporter is decreased [John & Jones 2007]. We therefore modelled SSRI effects by an increased value of *K*_*m*_ [Joshi *et al*. 2017].

**Figure 3:**
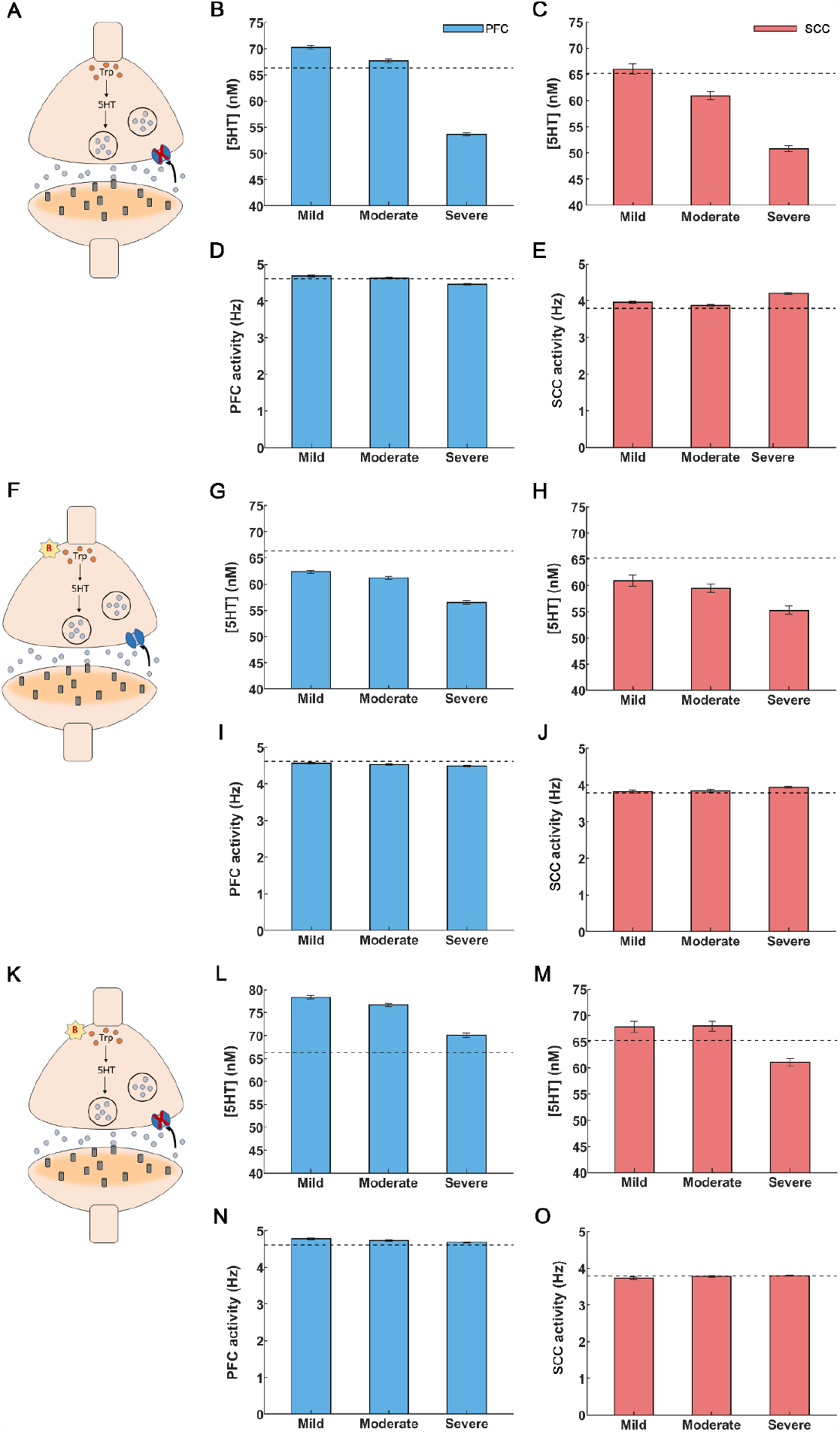
Drug interventions for inflammation-induced depression. **[A-E]** SSRI treatment. **[A]** A synapse under depression: serotonin synthesis is decreased and reuptake is increased, cf. (Figure 1A). SSRI application blocked serotonin transporters shown as a red “X”. **[B]** Administration of SSRIs led to a rise in PFC serotonin levels. SSRIs restored serotonin levels for mild and moderate inflammation but failed to do so for severe inflammation. **[C]** SCC serotonin was restored to control levels only for mild inflammation. **[D]** PFC activity was restored to control levels for mild and moderate inflammation. **[E]** Elevation in SCC activity was alleviated for mild and moderate inflammation. **[F-J]** Anti-inflammatory treatment. **[F]** Anti-inflammatory drugs caused a reduction in cytokines, depicted by a red “B” **[G**,**H]** Anti-inflammatory drugs raised serotonin levels but were insufficient to fully restore them to control levels for PFC and SCC, regardless of the degree of inflammation. **[I**,**J]** PFC and SCC activity were restored to control levels (PFC activity increased and SCC activity decreased). **[K-O]** Co-administration of SSRIs and anti-inflammatory treatment. **[K]** Synapse in depression undergoing simultaneous treatment with SSRIs (red “X”) and anti-inflammatory drugs (red “B”). **[L]** Extracellular serotonin levels were restored in the PFC for all inflammation conditions. **[M]** SCC serotonin levels were restored for mild and moderate inflammation. **[N**,**O]** Co-administration of SSRIs and anti-inflammatory drugs brought back PFC and SCC activity to control levels for all the inflammation conditions. The light blue (light red) bars in the figures depict extracellular serotonin and activity levels for the PFC (SCC) brain region. The black dashed lines refer to extracellular serotonin levels and activity in controls for PFC and SCC brain regions. The error bars refer to the sem values obtained from 100 simulations run for each condition.

We varied *K*_*m*_ between 170 (controls) to 200, similar to fast scan cyclic voltammetry experiments by [John & Jones 2007]. In that earlier work, serotonin concentration was measured in real time. The resulting estimates were fitted to the Michaelis-Menten based kinetic model and the constants *V*_*max*_ and *K*_*m*_ were obtained *(Methods)*. Our model predicted that SSRI administration increases PFC and SCC serotonin concentration (Figure 3B,C). For mild and moderate inflammation, PFC serotonin concentration was 70.26±0.36nM and 67.72±0.35nM respectively (Figure 3B, leftmost and middle bars). Interestingly, the latter value (moderate inflammation) is close to control levels, i.e. 66.35±0.35nM (shown by a dash line in Figure 3B, see also Figure 2C and [Hersey *et al*. 2021]).

However, for severe inflammation serotonin concentration was 53.63±0.26nM (Figure 3B, rightmost bar). In other words, SSRI administration failed to restore serotonin to control levels. In SCC, our model predicted a serotonin concentration of 66.01±0.93nM, 60.92±0.76nM and 50.80±0.55nM under mild, moderate and severe inflammation (Figure 3C). SSRIs restored serotonin levels to control levels (65.21±1.32nM, shown by a dashed line in Figure 3C) for mild inflammation only—not moderate nor severe. This is distinct from PFC, where SSRIs restore serotonin levels to control conditions, for mild and moderate inflammation.

We then assessed the impact of SSRIs on brain activity. For mild and moderate inflammation, SSRI administration resulted in an increased PFC activity of 4.68±0.02Hz and 4.63±0.02Hz respectively. These values are close to control levels (4.61±0.02Hz, [Kim & Sejnowski 2021; Kennedy *et al*. 2001], Figure 3D). Further, SSRI administration caused a reduction in SCC activity to 3.96±0.03Hz and 3.88±0.03Hz for mild and moderate inflammation. This is also similar to control activity (3.79±0.02Hz, Figure 3E; Kennedy *et al*. 2001). However, for severe inflammation SSRI treatment led to a PFC activity of 4.46±0.02Hz and SCC activity of 4.21±0.03Hz (Figures 3D,E, rightmost bars). Thus, SSRIs cannot restore brain activity to control levels in severe inflammation, which could be explained by the failure to restore serotonin concentration found above. This suggests that besides SSRIs, alternative treatments like targeting inflammation can be considered. This is what we did next. We studied anti-inflammatory drug effects on serotonin and brain activity.

### Anti-inflammatory drugs restore impaired brain activity but not serotonin deficiency

Above, we saw that SSRIs do not alleviate serotonin deficiency in the cingulo-frontal network under severe inflammation. This could be due to elevated cytokine levels that are not reduced by merely blocking serotonin reuptake. We thus asked if anti-inflammatory drugs could remedy serotonin deficiency and impaired brain activity.

In our model, anti-inflammatory drug administration was modelled through changing the value of the parameter *B* (*Methods*). This parameter characterizes the percentage reduction of cytokine concentration that affects both serotonin synthesis and reuptake (Equations 13 and 14 in *Methods*, red “B” in Figure 3F). We chose the parameter *B* (*B=*0.55), so that PFC serotonin levels rise to control levels when SSRIs and anti-inflammatory drugs are co-administered; as observed in [Hersey *et al*. 2021, Bai *et al*. 2020; Köhler-Forsberg *et al*. 2019]. This is discussed in the next section. Here, we consider the effect of anti-inflammatory drugs only. Our model predicted that for mild inflammation, anti-inflammatory drug administration increased cortical serotonin level to 62.35±0.32nM (leftmost bar in Figure 3G), away from control levels of 66.35±0.35nM (dash line in Figure 3G, [Hersey *et al*. 2021]). Further, for moderate and severe inflammation our model predicted serotonin levels to be 61.23±0.33nM and 56.53±0.29nM. Therefore, anti-inflammatory treatment caused an increase in serotonin levels but failed by itself to fully restore PFC serotonin to control levels regardless of inflammation degree. A similar response to anti-inflammatory drugs was found for SCC. Serotonin levels were only partially restored: 60.91±1.08nM, 59.45±0.83nM and 55.24±0.77nM for mild, moderate and severe inflammation respectively. These are also away from control levels, shown by a dash line in (Figure 3H).

The results for brain activity were different. Interestingly, administration of anti-inflammatory drugs restored impaired neural activity to control levels for all degrees of inflammation and in both brain areas. Estimates (bars) in Figures (Figures 3I,J) almost overlap with control levels (shown with dash line, Figures 3I,J). Specifically, following anti-inflammatory drugs administration PFC activity was restored to 4.56±0.02Hz (mild inflammation) and 4.53±0.02Hz (moderate). Similarly, SCC activity to 3.82±0.03Hz (mild) and 3.84±0.03Hz (moderate). For severe inflammation, the corresponding values were 4.49±0.02Hz (PFC, rightmost bar in Figure 3I) and 3.94±0.03Hz (SCC, rightmost bar in Figure 3J).

In brief, administering anti-inflammatory drugs did not restore serotonin concentration to control levels. It seems, however, that the new concentrations were able to change gating dynamics and input currents and restore brain activity. This could not be achieved using SSRIs. This also points towards combined pharmacological treatments, to which we turned next.

### Simultaneous SSRI and anti-inflammatory drug administration fully restored serotonin concentration to control levels

The foregoing numerical studies suggest that administering SSRIs or anti-inflammatory drugs on their own did not restore serotonin to control levels. Thus, we next simulated the effect of co-administering both drugs. We set the corresponding parameters in our model to treatment values, that is, *K*_*m*_ to 200 and *B* to 0.55. Recall that *K*_*m*_ reflects the rate of serotonin reuptake that is reduced by SSRI application. Further, *B* quantifies the percentage reduction in cytokine concentration as a result of anti-inflammatory drugs. Co-administering them led to an increase in serotonin concentration. Serotonin reuptake was reduced, and serotonin synthesis increased (red “B” and red “X” in Figure 3K). For severe inflammation, PFC serotonin levels were higher than control levels (66.35±0.35nM) and had a value equal to 70.07±0.39nM (Figure 3L, rightmost bar). Similarly, for SCC, the corresponding serotonin levels were 61.06±0.77nM similar to control levels of 65.21±1.32nM (Figure 3M, rightmost bar). Briefly, an increase in serotonin levels regardless of the severity of inflammation was observed. These results suggest that a joint treatment using SSRIs and anti-inflammatory drugs can remedy serotonin deficiency similar to observations by [Bai *et al*. 2020; Köhler-Forsberg *et al*. 2019; Brundin & Achtyes 2019].

Co-administering SSRIs and anti-inflammatory drugs also restored brain activity across all degrees of inflammation (Figures 3N,O). This could be driven by anti-inflammatory drugs that were found earlier to restore activity to control levels when administered on their own. In the case of dual administration, — for the case of severe inflammation — the PFC activity was 4.68±0.02Hz and SCC activity was 3.80±0.02Hz (Figures 3N,O, rightmost bars). These values were similar to control levels: 4.61±0.02Hz and 3.79±0.02Hz in PFC and SCC respectively (Wilson *et al*. 1994, Kim & Sejnowski 2021, Figures 2E,F, leftmost bar). In summary, we found that anti-inflammatory treatment together with SSRIs restored both network activity and neurotransmitter function.

In a separate set of analyses, we considered another effect of inflammation: besides serotonin, inflammation affects NMDA receptors and changes glutamate levels. This is due to degradation of tryptophan through the kynurenine pathway. Tryptophan degradation increases kynurenine metabolites [Dantzer & Walker 2014; Brydges *et al*. 2022]. One of them, quinolinic acid, is an NMDA receptor agonist that also increases extracellular glutamate [Müller & Schwarz 2007]. It blocks glutamate reuptake through astrocytes by reducing amino acid transporter 2 (EAAT2) [Miller & Raison 2016; Haroon *et al*. 2017]. This is important in depression, as elevated levels of quinolinic acid have been associated with inflammation and suicidal attempts [Dantzer & Walker 2014]. NMDA receptor activation of the sort induced by inflammation is known to play a role in depression. Depression symptoms were reduced following the administration of ketamine (an NMDA receptor antagonist) [Dantzer & Walker 2014]. Also, experimental findings by [Walker *et al*. 2013] demonstrated that inflammation-induced depression is mediated by NMDA receptors. To model these effects, we increased the NMDA time constant by 5% in SCC [Steiner *et al*. 2011], see also [Ramirez-Mahaluf *et al*. 2017; Alexander et al., 2019].

Separate administration of SSRIs and anti-inflammatory drugs did not restore control serotonin and brain activity levels. Our model predicted a reduction in serotonin and an increase in SCC activity after increasing the NMDA receptor time constant (Figures S1A,B). SSRI administration reversed these effects on serotonin in SCC but failed to restore control function (Figure S1C). SCC serotonin concentration was 63.34±0.82nM, 62.19±0.84nM and 50.15±0.56nM for mild, moderate and severe inflammation respectively (control levels were 65.21±1.32nM, shown using dash lines in Figure S1C). SCC activity was restored to control levels for mild inflammation (3.79±0.02Hz shown through dash lines in Figure S1D) but not for moderate and severe inflammation. Activity levels for mild, moderate and severe inflammation were 3.87±0.02Hz, 3.94±0.03Hz and 4.18±0.03Hz respectively.

Anti-inflammatory drugs in isolation also led to a serotonin increase but failed to achieve control concentrations (Figure S1E). SCC serotonin concentrations were found to be 61.41±0.89nM, 61.63±1.02nM and 55.02±0.77nM for mild, moderate and severe inflammation respectively. Further, SCC activity was restored to control levels (Figure S1F). It was 3.87±0.03Hz, 3.90±0.03Hz and 3.97±0.03Hz for mild, moderate and severe inflammation respectively. Similarly, to the results above, the control state was restored using a combined pharmacological treatment. Co-administration of SSRIs and anti-inflammatory drugs led to a complete restoration of serotonin concentrations: 74.84±1.28nM, 69.46±0.94nM and 63.55±0.86nM (Figure S1G). SCC activity levels were also restored: 3.91±0.03Hz, 3.82±0.03Hz and 3.91±0.02Hz.

### Reduced postsynaptic receptor density as a mechanism to cope with depression

The preceding analyses focused primarily on drug effects on presynaptic neurons. Newer drugs target postsynaptic neurons. Postsynaptic 5HT1A receptor density in SCC has been found to be reduced in depressed individuals [Wang *et al*. 2016; Shively *et al*. 2006]. Thus, in the last set of analyses, we used our model to predict changes in serotonin concentration and SCC activity for reduced postsynaptic receptor density.

We quantified postsynaptic receptor density by a parameter *R* and reduced this parameter by about 8% in accordance with [Meltzer *et al*. 2004]. This change left PFC serotonin levels unaffected (not shown). SCC serotonin levels for mild, moderate and severe inflammation were 66.64±0.96nM, 62.18±0.81nM and 49.16±0.48nM (Figure S2A). They were higher than the corresponding values of 56.63±0.82nM,54.16±0.66nM and 42.25±0.40nM when receptor changes were not included (Figure 2D). Thus, reducing postsynaptic receptor density could act as a compensatory mechanism used by the SCC to cope with serotonin deficiency. At the same time, SCC hyperactivity persisted. SCC activity was 4.40±0.03Hz, 4.38±0.02Hz and 4.66±0.03Hz for mild, moderate and severe inflammation respectively (Figure S2B).

We then asked what the effect of a simultaneous SSRI treatment would be. Following SSRI administration, serotonin rose above control levels for mild and moderate inflammation: 76.11±1.13nM and 73.35±1.02nM (control levels were 65.21±1.32nM, and are shown by dashed lines in Figure S1C). As with the previous results (that did not consider reduced postsynaptic receptor density, Figure 3B), this was not the case for severe inflammation. Our model predicted a serotonin concentration of 55.37±0.60nM for severe inflammation. The corresponding SCC activity levels were 4.39±0.03Hz, 4.36±0.03Hz and 4.50±0.03Hz, well above control levels with persisting hyperactivity (3.79±0.02Hz, dash line in Figure S2D).

On the other hand, application of anti-inflammatory drugs restored serotonin levels for mild and moderate inflammation (72.61±1.28nM and 68.88±1.10nM respectively, Figure S2E). This was not the case in our earlier results (Figure 3H). Overall, anti-inflammatory drugs alone could restore serotonin levels only in those individuals where postsynaptic receptor density was reduced and when inflammation was mild or moderate. This could explain experimental studies that found anti-inflammatory drugs to be effective in depression [Bai *et al*. 2020; Köhler *et al*. 2014].

The model also predicted SCC activity — under anti-inflammatory drugs – to be 4.27±0.03Hz, 4.24±0.03Hz and 4.35±0.03Hz for mild, moderate and severe inflammation, respectively (Figure S2F). Thus, neither SSRIs nor anti-inflammatory drugs could return activity back to control levels when administered separately. As before, this normalisation was seen only after combined drug administration. Then, serotonin concentration was restored to 71.53±1.06nM and activity to 4.21±0.03Hz for severe inflammation (Figures S2G,H) [Hersey *et al*. 2021]. For mild and moderate inflammation, serotonin concentration was 80.49±1.3nM and 78.27±1.24nM and the activity was 4.11±0.03Hz and 4.12±0.03Hz respectively.

## Discussion

We have presented a computational model that describes how inflammation changes extracellular serotonin levels and brain activity. Also, how brain activity, in turn, changes serotonin concentration, creating a feedback loop, leading to further changes in cortical and limbic activity, of the kind observed in depression.

Our model includes a simple, two—area cingulo-frontal circuit comprising PFC and SCC. Although these areas are not anatomically connected, they are known to be important in depression, show strong functional connectivity [Fox *et al*. 2012] and share a common DRN drive [Baumann *et al*. 2002]. Circuit dynamics are important in depression and its recurrence. One cause for recurrence is inflammation. This leads to changes in the HPA axis and alters cytokine levels in the cerebrospinal fluid [Köhler *et al*. 2017; Hiles *et al*. 2012; Hestad *et al*. 2003]. Cytokines are small signalling proteins that mediate communication between immune system cells and up-regulate inflammatory reactions. They are associated with multiple sclerosis and other conditions [Bonaccorso *et al*. 2001; Loftis & Hauser 2004]. Cytokine inhibition reduces depressive symptoms [Raison *et al*. 2013]. Here, we focused on the cytokine TNFα that is prevalent in depression (and inflammation) and described changes under different levels of TNFα concentration.

We considered changes for three levels of inflammation: mild, moderate and severe. This taxonomy was based on the severity of inflammation-induced depression reported by [Zou *et al*. 2018]. We formulated an index; namely, the degree of inflammation. This described inflammation severity. A higher degree of inflammation was associated with higher HAMD scores and disease severity.

Our model describes changes in brain activity for mild, moderate and severe inflammation and predicted the effects of SSRIs and anti-inflammatory drugs. For high degrees of inflammation, the model predicted a reduction in cortical (PFC) resting state activity and an increase in limbic (SCC) activity, similar to recordings by [Fernández-Palleiro *et al*. 2020; Wolff *et al*. 2019]. SCC hyperactivity — of the sort predicted here — has been used to identify responders [Mayberg *et al*. 1997]. We also found that SSRI action was limited by excessive cytokine concentration observed during severe inflammation [Hersey *et al*. 2021]. Our model predicted that only for mild inflammation can SSRIs alleviate depression. [Syed *et al*. 2018] also found lower cytokine levels in responders vs. non-responders to SSRI treatments. What distinguishes these two groups is still unclear. Our results suggest that this could be due to different levels of inflammation. Indeed, [Bhattacharya *et al*. 2019] found increased cytokine levels in non-responders after SSRI administration.

When either antidepressants or anti-inflammatory drugs were administered in isolation, our model predicted that serotonin concentration could be restored for mild and moderate inflammation only. To restore control levels of serotonin and brain activity in severe depression SSRIs and anti-inflammatory drugs needed to be co-administered simultaneously. This has also been observed in patients under a combined administration of anti-inflammatory drugs cyclooxygenase-2 (COX-2) inhibitors and SSRIs [Müller *et al*. 2006; Akhondzadeh *et al*. 2009]. Akhondzadeh *et al*. demonstrated the role of the anti-inflammatory agent, celecoxib, as an adjuvant with the SSRI fluoxetine. Over the six-week trial period, fluoxetine and celecoxib demonstrated superiority over fluoxetine administration alone. Similar results were obtained by Müller *et al*. who demonstrated that celecoxib, an anti-inflammatory drug, combined with the antidepressant reboxetine showed a significant reduction in HAMD scores. Animal studies [Casolini *et al*. 2002] also found that COX2 inhibition is associated with a reduced increase in the proinflammatory cytokines TNFα and IL1β together with lower anxiety and cognitive decline. Targeting cytokines and their signalling pathways has also been found to alleviate depression conditions [Dantzer *et al*. 2008; Wohleb *et al*. 2016; Raison *et al*. 2006].

Another inflammation effect is glutamate excitotoxicity [Miller *et al*. 2009; McNally *et al*. 2008; Müller & Schwarz 2007]. Activation of the kynurenine pathway activates NMDA receptors [McNally *et al*. 2008; Müller & Schwarz 2007] and contributes to excitotoxicity [Miller *et al*. 2009]. We considered these effects here. Similar to earlier results, a combined administration of SSRIs and anti-inflammatory drugs was needed to restore control serotonin and brain activity levels in severe depression.

Our modelling follows the work of [Ramirez-Mahaluf *et al*. 2017]. This work studied abnormalities in brain dynamics of depressed individuals as a result of glutamate metabolism in the ventral anterior cingulate cortex (vACC). A study by [Kringelbach *et al*., 2020] also introduced a biophysical model of serotonergic effects on brain activity. Here, we considered similar effects and their dependence on cytokine levels due to inflammation. Our model could be extended to include other brain regions that have been implicated in depression [Krishnan & Nestler 2008], including the amygdala, hippocampus, thalamus, striatum, and the parietal lobe [Zhang *et al*. 2017].

In previous work, we focused on changes in effective connectivity in depressed patients. We analysed data from a cognitive (MSIT) task that is commonly used to assess depressive state [Pinotsis *et al*. 2022]. We found that individual variability was explained by the feedforward drive from sensory areas to prefrontal cortex, alongside changes in neural activity in caudal areas. Connectivity changes best on synaptic plasticity. Here, we modelled such mechanisms in detail, in terms of changes in pre and postsynaptic receptor densities. A limitation of our model is that it cannot account for the 5HT1A autoreceptor changes but only captures postsynaptic receptors associated alterations. Selectively targeting autoreceptors can increase serotonergic neurotransmission and alleviate depression symptoms [Albert *et al*. 2011]. This acts as a brake and downregulates serotonin synthesis. This mechanism is thought to underlie delayed antidepressant response [Richardson-Jones *et al*. 2010] — and will be pursued under our model elsewhere.

In summary, elevated TNFα levels and inflammation left unabated, can lead to recurrence of depressive episodes [Miller & Raison 2016]. An understanding of the intricate links between the immune system and depression could prevent this remitting, relapsing trajectory. Our model is a first step in this direction. It describes the interaction between neurotransmission underlying depression and the immune system. It can model the combined effects of antidepressants and anti-inflammatory drugs and assesses their relevance with regard to depression severity. Further work is needed to understand white matter damage associated with inflammation that impairs neuronal communication [Pleasure *et al*. 2006] and how TMS and DBS affects neurotransmitter and immune systems. We hope that our work is a step towards a mechanistic understanding of depression, the role of inflammation and potential treatments.

## Methods

We studied the impact of peripheral inflammation on a cingulo-frontal network associated with depression. We used a Dynamic Mean Field model [Kringelbach *et al*. 2020]. This includes two subnetworks for the prefrontal cortex (PFC) and the subcallosal cingulate cortex (SCC). The activity of both these brain regions is impacted in depression. Inflammation was modelled here through elevated levels of the cytokine TNFα.

The population firing rates of the PFC and SCC subnetworks are given by

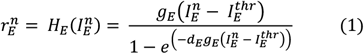

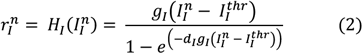

Here, *n* = {*PFC, SCC*} labels the PFC and SCC brain region. 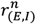 describes the excitatory (E) and inhibitory (I) neuronal population firing activity of the PFC and SCC brain regions. Within each brain region these neuronal populations are reciprocally connected to each other (Figure 1B, left panel). The neuronal response function *H*_(*E,I*)_ (Abbott & Chance 2005) serves as an input-output function transforming the currents 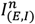 to produce *r*_(*E,I*)_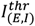 defines the threshold current. Further, the parameters *g*_(*E,I*)_ and *d*_(*E,I*)_ define the gain factor for the slope and curvature of *H*_(*E,I*)_ around 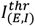 respectively.

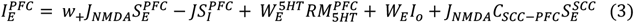

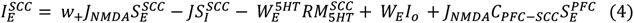

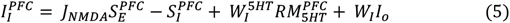

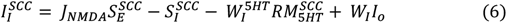

The currents 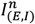 include inputs from excitatory and inhibitory neuronal populations, serotonergic currents 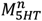, and external currents *I*_*o*_. *I*_*o*_ are scaled by the weights *W*_*E*_ and *W*_*I*_ for the E and I populations respectively. Further, *w*_+_ refers to the recurrent excitation weight, *J*_*NMDA*_ is the excitatory synaptic coupling and *J* is the local feedback synaptic coupling.

Both regions contain 5HT1A and 5HT2A receptors. PFC has an abundance of excitatory serotonergic 5HT2A receptors whereas the SCC is dominated by inhibitory serotonergic 5HT1A receptors [Palomero-Gallagher *et al*. 2009, Figure 1A,B]. For simplicity, we considered contributions from 5HT2A receptors in PFC and 5HT1A receptors in SCC. This is described by the parameter *R*. Also, 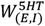 weight the excitatory and inhibitory inputs from the serotonergic system. PFC and SCC interact through bidirectional long-range excitatory connections (Figure 1C, right panel). *C*_*SCC*-*PFC*_ and *C*_*PFC*-*SCC*_ are the coupling constants between these brain regions. Synaptic dynamics are modelled by the following equations:

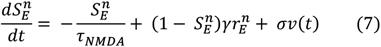

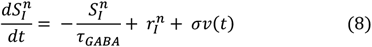

The synaptic gating variables, 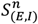 depend on the time constants of NMDA, *τ*_*NMDA*_, and GABA, *τ*_*GABA*_, respectively. *ν*(*t*) is the uncorrelated standard Gaussian noise with an amplitude *σ*. Serotonergic current, 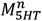, dynamics are given by

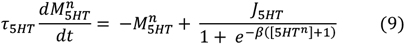

The serotonin concentration appearing above changes over time as follows:

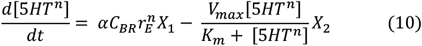

where *X*_1_ and *X*_2_ are given by

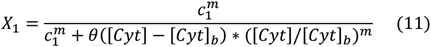

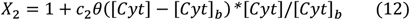

Below we explain the parameters appearing above. Serotonin concentration [5*HT*^*n*^], and its dynamics following synthesis and reuptake are modelled by Equation 10. This is a Michaelis-Menten kinetic scheme, where *α* controls the serotonergic current so that the drive in Equation 9 is around the centre of the sigmoid. *C*_*BR*_ is the fiber density connectivity between the prefrontal cortex (SCC) and the raphe nuclei. *V*_*max*_ and *K*_*m*_ are the Michaelis-Menten constants that define the maximum re-uptake rate and the serotonin concentration at which the re-uptake rate is half of the maximum rate respectively. The expression [*Cyt*]/[*Cyt*]_*b*_ is the degree of inflammation. [*Cyt*]_*b*_ is the basal TNFα cytokine concentration in controls and [*Cyt*] is the elevated TNFα cytokine level under inflammation. The degree of inflammation takes only positive values, see Results and [Zou *et al*. 2018] for details. The term *X*_1_ in Equation 10 models the cytokine effects on serotonin synthesis as a result of reduced tryptophan. *c*_1_ and *m* defines the shape and steepness of *X*_1_. The term *X*_2_ in Equation 10 describes the rapid increase in serotonin re-uptake. *c*_2_ describes the increased re-uptake rate. *τ*_5*HT*_, *J*_5*HT*_ and *β* are the parameters that define the time constant, range and slope of the serotonergic currents.

### Modelling drug treatments

In this study, various drug treatments were modelled by the following parameter changes.

1. **SSRI treatment:** The value of *K*_*m*_ in Equation 10 is varied from 170nM to 200nM.
2. **Anti-inflammatory treatment:** The parameter *B* in Equations 13 and 14. models vs. anti-inflammatory blocker effects (*B* =0.55)

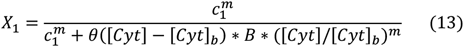

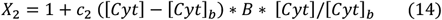
3. **Co-administration of SSRIs and anti-inflammatory drug treatment:** Dual treatment is implemented by simultaneously changing *K*_*m*_ to 200nM and *B* to 0.55. All simulations were performed using MATLAB. The differential equations were solved numerically using the Euler-Maruyama method with a time step of 0.1ms. Simulations ran for 7 seconds of simulated time. We ran 100 simulations for each condition. This data was then used to obtain the mean and sem values. The values of the various parameters used in the simulations are provided in Tables 1 and 2.

## Acknowledgements

This work was supported by the Economic and Social Research Council (ESRC) (Grant Number ES/T01279X/1).

## Footnotes

1 Note that these authors report measurements in mice. Because concentrations are normalized to unit volume, we assumed that changes from the control condition for different levels of inflammation will be similar across species. To produce inflammation and induce depression-like behaviour Hersey *et al*. [Hersey *et al*. 2021] administered lipopolysaccharide (LPS), a known immune system stimulator.

## Notes

### Competing Interest Statement

The authors have declared no competing interest.

## References

1. Depression and other common mental disorders: global health estimates. Geneva: World Health Organization; 2017. Licence: CC BY-NC-SA 3.0 IGO

2. Malhi GS, Mann JJ (2018) Depression. Lancet 392: 2299–2312

3. Sarter M, Markowitsch HJ (1984) Collateral innervation of the medial and lateral prefrontal cortex by amygdaloid, thalamic, and brain-stem neurons. J Comp Neurol 224:445–460

4. Mayberg HS (2007) Defining the neural circuitry of depression: toward a new nosology with therapeutic implications. Biol Psychiatry 61:729–730

5. Dunlop BW, Cha J, Choi KS, Rajendra JK, Nemeroff CB, Craighead WE, Mayberg HS (2023) Shared and unique changes in brain connectivity among depressed patients after remission with pharmacotherapy versus psychotherapy. Am J Psychiatry 180: 218–229

6. Gray JP, Manuello J, Alexander-Bloch AF, Leonardo C, Franklin C, Choi KS, Cauda F, Costa T, Blangero J, Glahn DC, Mayberg HS, Fox PT (2023) Co-alteration network architecture of major depressive disorder: A multi-modal neuroimaging assessment of large-scale disease effects. Neuroinform 21: 443–455

7. Mayberg HS, Brannan SK, Mahurin RK, Jerabek PA, Brickman JS, Tekell JL, Silva JA, McGinnis S, Glass TG, Martin CC, Fox PT (1997) Cingulate function in depression a potential predictor of treatment response. Neuroreport 8:1057–1061

8. Mayberg HS, Liotti M, Brannan SK, McGinnis S, Mahurin RK, Jerabek PA, Arturo Silva J, Tekell JL, Martin CC, Lancaster JL, Fox PT (1999) Reciprocal limbic-cortical function and negative mood: Converging PET findings in depression and normal sadness. Am J Psychiatry 156:675–682

9. Greicius MD, Flores BH, Menon V, Glover GH, Solvason HB, Kenna H, Reiss AL, Schatzberg AF (2007) Resting-State functional connectivity in major depression: Abnormally increased contributions from subgenual cingulate cortex and thalamus. Biological Psychiatry 62: 429–437.

10. Liu Y, Chen Y, Liang X, Li D, Zheng Y, Zhang H, Cui Y, Chen J, Liu J and Qiu S (2020) Altered resting-state functional connectivity of multiple networks and disrupted correlation with executive function in major depressive disorder. Front Neurol 11:272

11. Touya M, Lawrence DF, Kangethe A, Chrones L, Evangelatos T, Polson M (2022) Incremental burden of relapse in patients with major depressive disorder: a real-world, retrospective cohort study using claims data. BMC Psychiatry 22: 152

12. Kessing LV, Andersen PK (2017) Evidence for clinical progression of unipolar and bipolar disorders. Acta Psychiatr Scand 135: 51–64

13. Gabriel FC, de Melo DO, Fráguas R, Leite-Santos NC, Mantovani da Silva RA, Ribeiro E (2020) Pharmacological treatment of depression: A systematic review comparing clinical practice guideline recommendations. PLoS one 15: e0231700

14. Voineskos D, Daskalakis ZJ, Blumberger DM (2020) Management of treatment-resistant depression: challenges and strategies. Neuropsychiatr Dis Treat 16:221–234

15. Miller AH, Maletic V, Raison CL (2009) Inflammation and its discontents: the role of cytokines in the pathophysiology of major depression. Biol Psychiatry 65: 732–741

16. Miller AH, Timmie WP (2009) Norman cousins lecture. Mechanisms of cytokine-induced behavioral changes: Psychoneuroimmunology at the translational interface. Brain Behav Immun 23: 149–158

17. Dantzer R, O’Connor JC, Freund GG, Johnson RW, Kelley KW (2008) From inflammation to sickness and depression: when the immune system subjugates the brain. Nat Rev Neurosci 9: 46–56

18. McEwen BS, Gianaros PJ (2011) Stress-and allostasis-induced brain plasticity. Annu Rev Med 62:431–445

19. Peters A, McEwen BS, Friston K (2017). Uncertainty and stress: Why it causes diseases and how it is mastered by the brain. Prog Neurobiol 156:164–188

20. McEwen BS (1998) Protective and damaging effects of stress mediators. N Engl J Med 338: 171–179

21. Cohen S, Janicki-Deverts D, Miller GE (2007) Psychological stress and disease. JAMA 298:1685–1688

22. Osimo EF, Baxter LJ, Lewis G, Jones PB, Khandaker GM (2019) Prevalence of low-grade inflammation in depression: a systematic review and meta-analysis of CRP levels. Psychol Med 49: 1958–1970

23. Carvalho LA, Torre JP, Papadopoulos AS, Poon L, Juruena MF, Markopoulou K, Cleare AJ, Pariante CM (2013) Lack of clinical therapeutic benefit of antidepressants is associated overall activation of the inflammatory system. J Affect Disord 148:136–140.

24. Yoshimura R, Hori H, Ikenouchi-Sugita A, Umene-Nakano W, Ueda N, Nakamura J (2009) Higher plasma interleukin-6 (IL-6) level is associated with SSRI-or SNRI-refractory depression. Prog Neuropsychopharmacol Biol Psychiatry 33:722–726

25. Hodes GE, Kana V, Menard C, Merad M, Russo SJ (2015) Neuroimmune mechanisms of depression. Nat Neurosci 18: 1386–1393

26. Page CE, Coutellier L (2019) Prefrontal excitatory/inhibitory balance in stress and emotional disorders: Evidence for over-inhibition. Neurosci Biobehav Rev 105: 39–51

27. Kringelbach ML, Cruzat J, Cabral J, Knudsen GM, Carhart-Harris R, Whybrow PC, Logothetis NK, Deco G (2020) Dynamic coupling of whole-brain neuronal and neurotransmitter systems. Proc Natl Acad Sci USA 117: 9566–9576

28. Ramirez-Mahaluf JP, Roxin A, Mayberg HS, Compte A (2017) A computational model of major depression: the role of glutamate dysfunction on cingulo-frontal network dynamics. Cereb Cortex 27: 660–679

29. Pariante CM, Lightman SL (2008) The HPA axis in major depression: classical theories and new developments. Trends Neurosci 31: 464–468

30. Bhat A, Parr T, Ramstead M, Friston K (2021) Immunoceptive inference: why are psychiatric disorders and immune responses intertwined? Biol Philos 36: 27

31. Raison CL, Rutherford RE, Woolwine BJ, Shuo C, Schettler P, Drake DF, Haroon E, Miller AH (2013) A randomized controlled trial of the tumor necrosis factor antagonist infliximab for treatment-resistant depression: the role of baseline inflammatory biomarkers. JAMA Psychiatry 70:31–41

32. Yao L, Pan L, Qian M, Sun W, Gu C, Chen L, Tang X, Hu Y, Xu L, Wei Y, Hui L, Liu X, Wang J, Zhang T (2020) Tumor necrosis factor-α variations in patients with major depressive disorder before and after antidepressant treatment. Front Psychiatry 11:518837

33. Brymer KJ, Romay-Tallon R, Allen J, Caruncho HJ, Kalynchuk LE (2019) Exploring the potential antidepressant mechanisms of TNFα antagonists. Front Neurosci 13:98

34. Dowlati Y, Herrmann N, Swardfager W, Liu H, Sham L, Reim EK, Lanctôt KL (2010) A metaanalysis of cytokines in major depression. Biol Psychiatry 67:446–457

35. Köhler CA, Freitas TH, Maes M, de Andrade NQ, Liu CS, Fernandes BS, Stubbs B, Solmi M, Veronese N, Herrmann N, Raison CL, Miller BJ, Lanctôt KL, Carvalho AF (2017) Peripheral cytokine and chemokine alterations in depression: a meta-analysis of 82 studies. Acta Psychiatr Scand 135: 373–387

36. Fox MD, Buckner RL, White MP, Greicius MD, Pascual-Leone A (2012) Efficacy of transcranial magnetic stimulation targets for depression is related to intrinsic functional connectivity with the subgenual cingulate. Biol Psychiatry 72: 595–603.

37. Zou W, Feng R, Yang Y (2018) Changes in the serum levels of inflammatory cytokines in antidepressant drug-naïve patients with major depression. PLoS ONE 13: e0197267.

38. Liu JJ, Wei YB, Strawbridge R, Bao Y, Chang S, Shi L, Que J, Gadad BS, Trivedi MH, Kelsoe JR, Lu L (2020) Peripheral cytokine levels and response to antidepressant treatment in depression: a systematic review and meta-analysis. Mol Psychiatry 25:339–350

39. Miller AH, Haroon E, Raison CL, Felger JC (2013) Cytokine targets in the brain: impact on neurotransmitters and neurocircuits. Depress Anxiety 30: 297–306.

40. Ma K, Zhang H, Baloch Z (2016) Pathogenetic and therapeutic applications of tumor necrosis factor-α (TNF-α) in major depressive disorder: a systematic review. Int J Mol Sci 17: 733

41. Hestad KA, Tønseth S, Støen CD, Ueland T, Aukrust P (2003) Raised plasma levels of tumor necrosis factor alpha in patients with depression: normalization during electroconvulsive therapy. J ECT 19:183–188.

42. Tyring S, Gottlieb A, Papp K, Gordon K, Leonardi C, Wang A, Lalla D, Woolley M, Jahreis A, Zitnik R, Cella D, Krishnan R (2006) Etanercept and clinical outcomes, fatigue, and depression in psoriasis: double-blind placebo-controlled randomised phase III trial. Lancet 367:29–35

43. Monk JP, Phillips G, Waite R, Kuhn J, Schaaf LJ, Otterson GA, Guttridge D, Rhoades C, Shah M, Criswell T, Caligiuri MA, Villalona-Calero MA (2006) Assessment of tumor necrosis factor alpha blockade as an intervention to improve tolerability of dose-intensive chemotherapy in cancer patients. J Clin Oncol 24:1852–1859

44. Persoons P, Vermeire S, Demyttenaere K, Fischler B, Vandenberghe J, Van Oudenhove L, Pierik M, Hlavaty T, Van Assche G, Noman M, Rutgeerts P (2005) The impact of major depressive disorder on the short- and long-term outcome of Crohn’s disease treatment with infliximab. Aliment Pharmacol Ther 22:101–110

45. Silverman MN, Macdougall MG, Hu F, Pace TW, Raison CL, Miller AH (2007) Endogenous glucocorticoids protect against TNF-alpha-induced increases in anxiety-like behavior in virally infected mice. Mol Psychiatry 12:408–417

46. Mulders PC, van Eijndhoven PF, Schene AH, Beckmann CF, Tendolkar I (2015) Resting-state functional connectivity in major depressive disorder: a review. Neurosci Biobehav Rev 56:330–344

47. Liston C, Chen AC, Zebley BD, Drysdale AT, Gordon R, Leuchter B, Voss HU, Casey BJ, Etkin A, Dubin MJ (2014) Default mode network mechanisms of transcranial magnetic stimulation in depression. Biol Psychiatry 76: 517–526

48. Froudist-Walsh S, Xu T, Niu M, Rapan L, Zhao L, Margulies DS, Zilles K, Wang X-J, Palomero-Gallagher N (2023) Gradients of neurotransmitter receptor expression in the macaque cortex. Nat Neurosci 26: 1281–1294

49. Palomero-Gallagher N, Vogt BA, Schleicher A, Mayberg HS, Zilles K (2009) Receptor architecture of human cingulate cortex: evaluation of the four-region neurobiological model. Hum Brain Mapp 30:2336–2355

50. Hersey M, Samaranayake S, Berger SN, Tavakoli N, Mena S, Nijhout HF, Reed MC, Best J, Blakely RD, Reagan LP, Hashemi P (2021) Inflammation-induced histamine impairs the capacity of escitalopram to increase hippocampal extracellular serotonin. J Neurosci 41: 6564–6577

51. Dum RP, Levinthal DJ, Strick PL (2016) Motor, cognitive, and affective areas of the cerebral cortex influence the adrenal medulla. Proc Natl Acad Sci USA 113:9922–9927

52. Wilson FAW, O’Scalaidhe SP, Goldman-Rakic PS (1994) Functional synergism between putative γ-aminobutyrate-containing neurons and pyramidal neurons in prefrontal cortex. Proc Natl Acad Sci USA 91:4009–4013

53. Kim R, Sejnowski TJ (2021) Strong inhibitory signaling underlies stable temporal dynamics and working memory in spiking neural networks. Nat Neurosci 24: 129–139

54. Fernández-Palleiro P, Rivera-Baltanás T, Rodrigues-Amorim D, Fernández-Gil S, Vallejo-Curto MDC, Álvarez-Ariza M, López M, Rodriguez-Jamardo C, Benavente JL, de Las Heras E, Olivares JM, Spuch C (2020) Brainwaves oscillations as a potential biomarker for major depression disorder risk. Clin EEG Neurosci 51: 3–9

55. Fitzgerald PB, Laird AR, Maller J, Daskalakis ZJ (2008) A meta-analytic study of changes in brain activation in depression. Hum Brain Mapp 29:683–695

56. Pizzagalli DA (2011) Frontocingulate dysfunction in depression: toward biomarkers of treatment response. Neuropsychopharmacology Reviews 36: 183–206

57. Mayberg HS, Lozano AM, Voon V, McNeely HE, Seminowicz D, Hamani C, Schwalb JM, Kennedy SH (2005) Deep brain stimulation for treatment-resistant depression. Neuron 45: 651–660

58. Wolff A, de la Salle S, Sorgini A, Lynn E, Blier P, Knott V and Northoff G (2019) Atypical temporal dynamics of resting state shapes stimulus-evoked activity in depression—An EEG study on rest–stimulus interaction. Front Psychiatry 10:719

59. Shao J, Meng C, Tahmasian M, Brandl F, Yang Q, Luo G, Luo C, Yao D, Gao L, Riedl V, Wohlschläger A, Sorg C (2018) Common and distinct changes of default mode and salience network in schizophrenia and major depression. Brain Imaging Behav 12: 1708–1719

60. John CE, Jones SR (2007) Voltammetric characterization of the effect of monoamine uptake inhibitors and releasers on dopamine and serotonin uptake in mouse caudate-putamen and substantia nigra slices. Neuropharmacology 52:1596–1605

61. Joshi A, Youssofzadeh V, Vemana V, McGinnity TM, Prasad G, Wong-Lin KF (2017) An integrated modelling framework for neural circuits with multiple neuromodulators. J R Soc Interface 14: 20160902

62. Kennedy SH, Evans KR, Krüger S, Mayberg HS, Meyer JH, McCann S, Arifuzzman AI, Houle S, Vaccarino F (2001) Changes in regional brain glucose metabolism measured with positron emission tomography after paroxetine treatment of major depression. Am J Psychiatry 158:899–905

63. Bai S, Guo W, Feng Y, Deng H, Li G, Nie H, Guo G, Yu H, Ma Y, Wang J, Chen S, Jing J, Yang J, Tang Y, Tang Z (2020) Efficacy and safety of anti-inflammatory agents for the treatment of major depressive disorder: a systematic review and meta-analysis of randomised controlled trials. J Neurol Neurosurg Psychiatry 91:21–32

64. Köhler-Forsberg O, Lydholm CN, Hjorthøj C, Nordentoft M, Mors O, Benros ME (2019) Efficacy of anti-inflammatory treatment on major depressive disorder or depressive symptoms: meta-analysis of clinical trials. Acta Psychiatr Scand 139:404–419

65. Brundin L, Achtyes E (2019) Has the time come to treat depression with anti-inflammatory medication? Acta Psychiatr Scand 139: 401–403

66. Dantzer R, Walker AK (2014) Is there a role for glutamate-mediated excitotoxicity in inflammation-induced depression? J Neural Transm (Vienna) 121: 925–932

66. Brydges CR, Bhattacharyya S, Dehkordi SM, Milaneschi Y, Penninx B, Jansen R, Kristal BS, Han X, Arnold M, Kastenmüller G, Bekhbat M, Mayberg HS, Craighead WE, Rush AJ, Fiehn O, Dunlop BW, Kaddurah-Daouk R (2022) Mood Disorders Precision Medicine Consortium. Metabolomic and inflammatory signatures of symptom dimensions in major depression. Brain Behav Immun 102: 42–52

67. Müller N, Schwarz MJ (2007) The immune-mediated alteration of serotonin and glutamate: towards an integrated view of depression. Mol Psychiatry 12: 988–1000

68. Miller AH, Raison CL (2016) The role of inflammation in depression: from evolutionary imperative to modern treatment target. Nat Rev Immunol 16: 22–34

69. Haroon E, Miller AH, Sanacora G (2017) Inflammation, glutamate, and glia: A trio of trouble in mood disorders. Neuropsychopharmacology 42:193–215

70. Walker AK, Budac DP, Bisulco S, Lee AW, Smith RA, Beenders B, Kelley KW, Dantzer R (2013) NMDA receptor blockade by ketamine abrogates lipopolysaccharide-induced depressive-like behavior in C57BL/6J mice. Neuropsychopharmacology 38:1609 –1616

71. Steiner J, Walter M, Gos T, Guillemin GJ, Bernstein H-G, Sarnyai Z, Mawrin C, Brisch R, Bielau H, zu Schwabedissen LM, Bogerts B, Myint A-M (2011) Severe depression is associated with increased microglial quinolinic acid in subregions of the anterior cingulate gyrus: evidence for an immune-modulated glutamatergic neurotransmission? J Neuroinflammation 8:94

72. Alexander L, Gaskin P L, Sawiak S J, Fryer TD, Hong YT, Cockcroft GJ, Clarke HF, Roberts AC (2019). Fractionating blunted reward processing characteristic of anhedonia by overactivating primate subgenual anterior cingulate cortex. Neuron 101: 307–320

73. Wang L, Zhou C, Zhu D, Wang X, Fang L, Zhong J, Mao Q, Sun L, Gong X, Xia J, Lian B, Xie P (2016) Serotonin-1A receptor alterations in depression: a meta-analysis of molecular imaging studies. BMC Psychiatry 16:319

74. Shively CA, Friedman DP, Gage HD, Bounds MC, Brown-Proctor C, Blair JB, Henderson JA, Smith MA, Buchheimer N (2006) Behavioral depression and positron emission tomography-determined serotonin 1A receptor binding potential in cynomolgus monkeys. Arch Gen Psychiatry 63: 396–403

75. Meltzer CC, Price JC, Mathis CA, Butters MA, Ziolko SK, Moses-Kolko E, Mazumdar S, Mulsant BH, Houck PR, Lopresti BJ, Weissfeld LA, Reynolds CF (2004) Serotonin 1A receptor binding and treatment response in late-life depression. Neuropsychopharmacology 29: 2258–2265

76. Köhler O, Benros ME, Nordentoft M, Farkouh ME, Iyengar RL, Mors O, Krogh J (2014) Effect of anti-inflammatory treatment on depression, depressive symptoms, and adverse effects: a systematic review and meta-analysis of randomized clinical trials. JAMA Psychiatry. 71:1381–1391

77. Baumann B, Bielau H, Krell D, Agelink MW, Diekmann S, Wurthmann C, Trübner K, Bernstein HG, Danos P, Bogerts B (2002) Circumscribed numerical deficit of dorsal raphe neurons in mood disorders. Psychol Med 32:93–103

78. Hiles SA, Baker AL, de Malmanche T, Attia J (2012) A meta-analysis of differences in IL-6 and IL-10 between people with and without depression: exploring the causes of heterogeneity. Brain Behav Immun 26:1180–1188

79. Bonaccorso S, Puzella A, Marino V, Pasquini M, Biondi M, Artini M, Almerighi C, Levrero M, Egyed B, Bosmans E, Meltzer HY, Maes M (2001) Immunotherapy with interferon-alpha in patients affected by chronic hepatitis C induces an intercorrelated stimulation of the cytokine network and an increase in depressive and anxiety symptoms. Psychiatry Res 105:45–55

80. Loftis JM, Hauser P (2004) The phenomenology and treatment of interferon-induced depression. J Affect Disord 82:175–190

81. Syed SA, Beurel E, Loewenstein DA, Lowell JA, Craighead WE, Dunlop BW, Mayberg HS, Dhabhar F, Dietrich WD, Keane RW, de Rivero Vaccari JP, Nemeroff CB (2018) Defective inflammatory pathways in never-treated depressed patients are associated with poor treatment response. Neuron 99: 914–924.e3

82. Bhattacharyya S, Dunlop BW, Mahmoudiandehkordi S, Ahmed AT, Louie G, Frye MA, Weinshilboum RM, Krishnan RR, Rush AJ, Mayberg HS, Craighead WE, Kaddurah-Daouk R (2019) Pilot study of metabolomic clusters as state markers of major depression and outcomes to CBT treatment. Front Neurosci 13:926

83. Müller N, Schwarz MJ, Dehning S, Douhe A, Cerovecki A, Goldstein-Müller B, Spellmann I, Hetzel G, Maino K, Kleindienst N, Möller H-J, Arolt V, Riedel M (2006) The cyclooxygenase-2 inhibitor celecoxib has therapeutic effects in major depression: results of a double-blind, randomized, placebo controlled, add-on pilot study to reboxetine. Mol Psychiatry 11: 680–684

84. Akhondzadeh S, Jafari S, Raisi F, Nasehi AA, Ghoreishi A, Salehi B, Mohebbi-Rasa S, Raznahan M, Kamalipour A (2009) Clinical trial of adjunctive celecoxib treatment in patients with major depression: a double blind and placebo controlled trial. Depress Anxiety 26: 607–611

85. Casolini P, Catalani A, Zuena AR, Angelucci L (2002) Inhibition of COX-2 reduces the age-dependent increase of hippocampal inflammatory markers, corticosterone secretion, and behavioral impairments in the rat. J Neurosci Res 68: 337–343

86. Wohleb ES, Franklin T, Iwata M, Duman RS (2016) Integrating neuroimmune systems in the neurobiology of depression. Nat Rev Neurosci 17: 497–511

87. Raison CL, Capuron L, Miller AH (2006) Cytokines sing the blues: inflammation and the pathogenesis of depression. Trends Immunol 27: 24–31

88. McNally L, Bhagwagar Z, Hannestad J (2008) Inflammation, glutamate, and glia in depression: a literature review. CNS Spectr 13:501–510.

89. Krishnan V, Nestler EJ (2008) The molecular neurobiology of depression. Nature 455: 894–902

90. Zhang F-F, Peng W, Sweeney JA, Jia Z-Y, Gong Q-Y (2018) Brain structure alterations in depression: Psychoradiological evidence. CNS Neurosci Ther 24: 994–1003

91. Pinotsis DA, Fitzgerald S, See C, Sementsova A and Widge AS (2022) Toward biophysical markers of depression vulnerability. Front Psychiatry 13:938694

92. Albert PR, L. François B, Millar AM (2011) Transcriptional dysregulation of 5-HT1A autoreceptors in mental illness. Mol Brain 4:21

93. Richardson-Jones JW, Craige CP, Guiard BP, Stephen A, Metzger KL, Kung HF, Gardier AM, Dranovsky A, David DJ, Beck SG, Hen R, Leonardo ED (2010) 5-HT1A autoreceptor levels determine vulnerability to stress and response to antidepressants. Neuron 65: 40–52

94. Pleasure D, Soulika A, Singh S K, Gallo V, Bannerman P (2006) Inflammation in white matter: clinical and pathophysiological aspects. Ment Retard Dev Disabil Res Rev 12: 141–146

95. Abbott LF, Chance FS (2005) Drivers and modulators from push-pull and balanced synaptic input. Prog Brain Res 149: 147–155

