## Supplementary material for "A computational account of joint SSRI and anti-inflammatory treatment": Figure S1, Figure S2: Supplementary Material - Reneaux et al. 2023.pdf

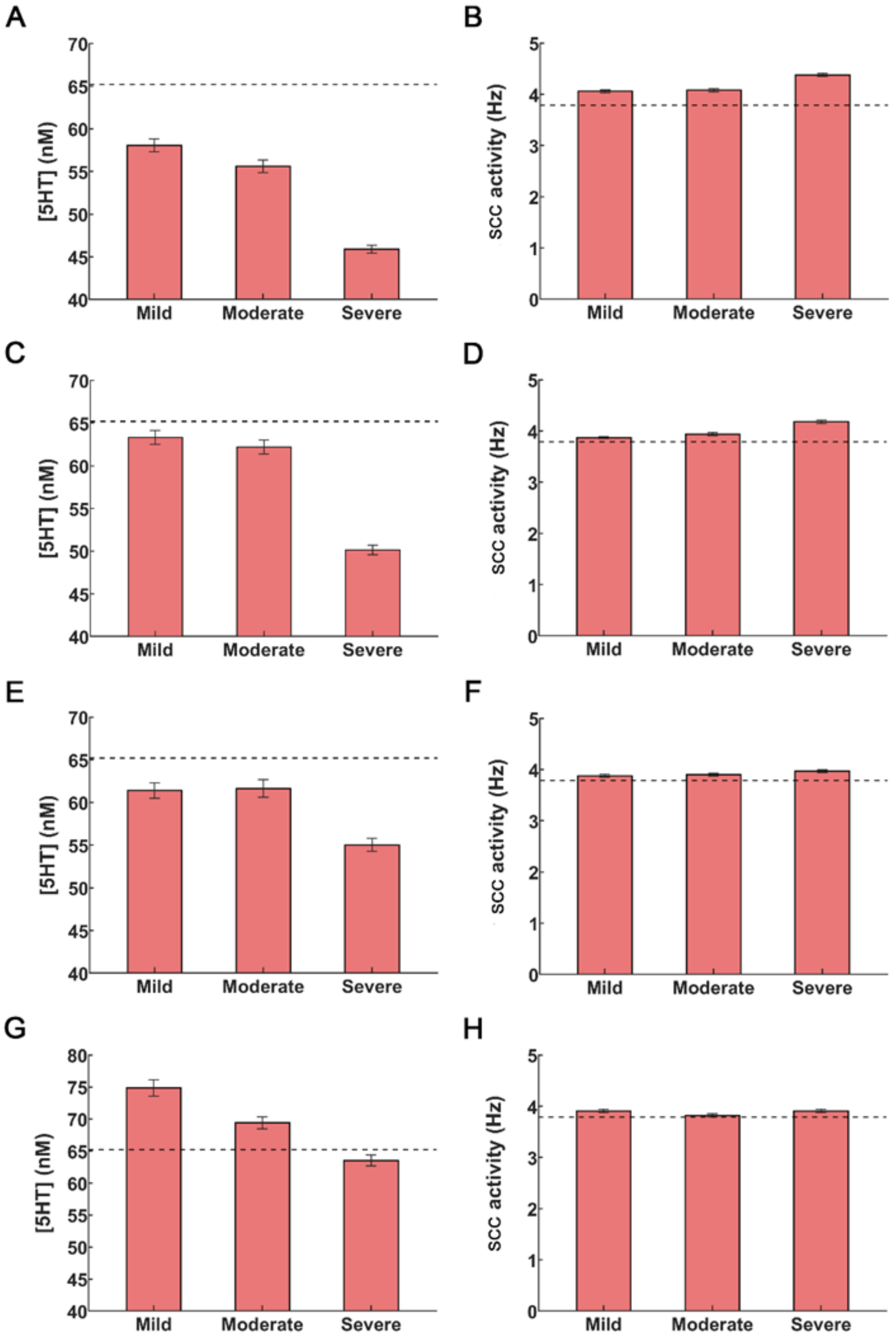


**Figure S1: Impact of NMDAR agonists on serotonin and SCC activity. [A]** Reduction in extracellular serotonin. **[B]** Increase in SCC activity. **[C]** Application of SSRIs increases SSC serotonin but fails to restore it. **[D]** Administration of SSRIs restores SCC activity for mild inflammation but not moderate and severe inflammation. **[E]** Anti-inflammatory drugs cause an elevation of serotonin levels but not to control levels.

**[F]** SCC activity for mild and moderate inflammation is restored **[G]** Co-administration of SSRIs and anti-inflammatory drugs restore serotonin to control levels **[H]** Co-administration of SSRIs and anti-inflammatory drugs also restores SCC activity. The black dash lines show control levels.

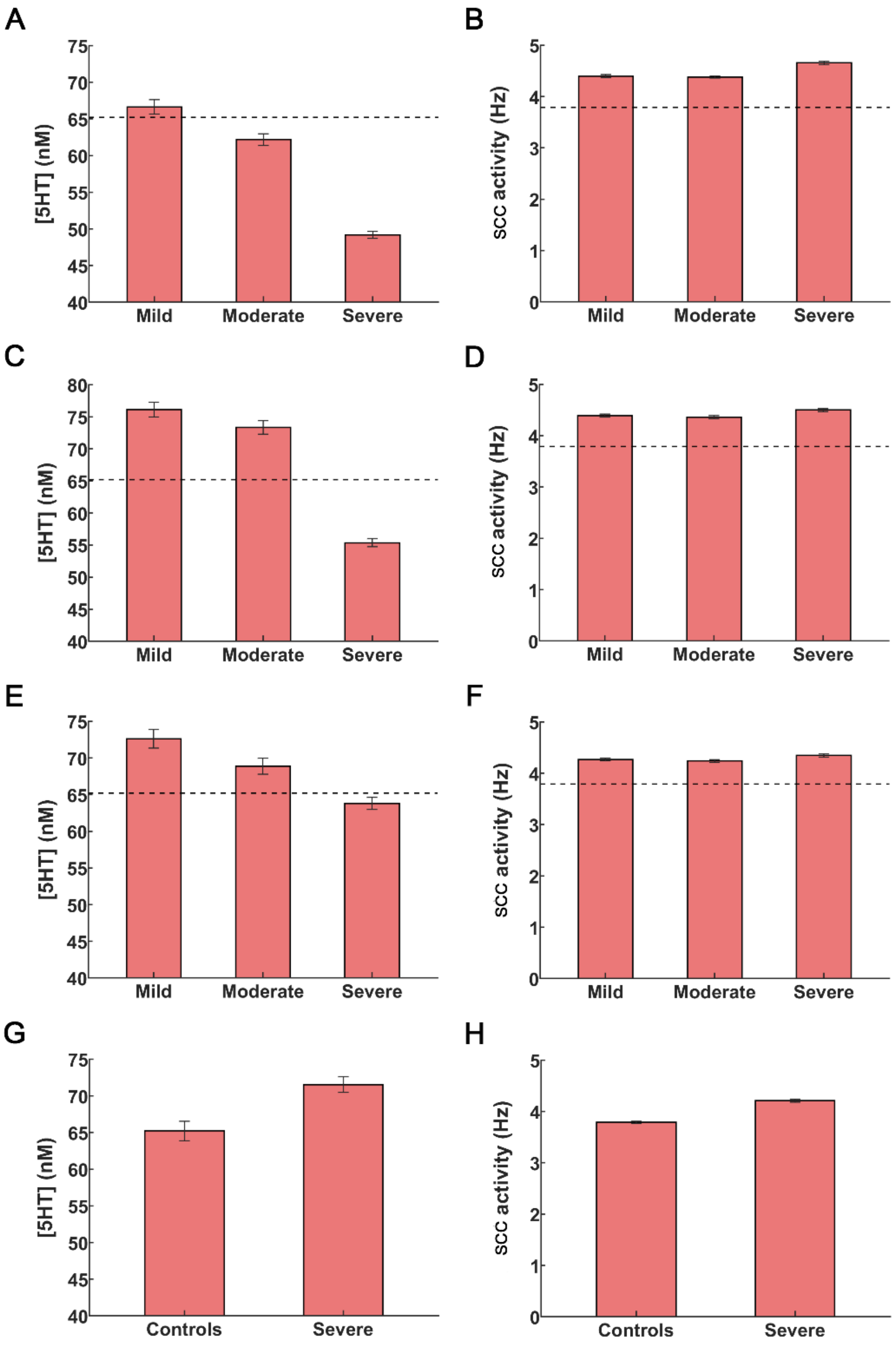


**Figure S2: Alterations in postsynaptic inhibitory serotonin receptors in the SCC as the brain's compensatory mechanism to combat depression. [A]** Reduction in 5HT1A receptor density restores SCC serotonin levels for mild inflammation and partially elevates serotonin levels for moderate and severe inflammation. **[B]** Elevated SCC activity above control levels persists for all the conditions of inflammation. **[C]** Application of SSRIs restores serotonin above control levels for mild and moderate inflammation. **[D]** SSRIs fail to reduce SCC hyperactivity to control levels. **[E]** Application of anti-inflammatory drugs restores SCC serotonin concentration for mild and moderate inflammation, see also (Figure 3H). **[F]** SCC activity under anti-inflammatory drug administration remains elevated across all degrees of inflammation. **[G,H]** Co-administration of SSRIs and anti-inflammatory drugs restores SCC serotonin levels for the severe inflammation condition. Isolated treatments with SSRIs (Figures 3C,E) or anti-inflammatory drugs (Figures 3H,J) were unable to achieve this. The dash line in the figure panels refers to control levels of serotonin concentration and activity in the SCC. Here, we have not shown the PFC serotonin and activity levels as they are very similar to the levels reported in Figure 3. The error bars refer to the sem values obtained from 100 simulations run for each condition.
